# Vesicle-associated *Pseudomonas aeruginosa* leucine aminopeptidase modulates bacterial biofilms grown on host cellular substrates

**DOI:** 10.1101/587279

**Authors:** Caitlin N. Esoda, Meta J. Kuehn

## Abstract

*Pseudomonas aeruginosa*, known as one of the leading causes of morbidity and mortality in cystic fibrosis (CF) patients, secretes a variety of virulence-associated proteases. These enzymes have been shown to contribute significantly to *P. aeruginosa* pathogenesis and biofilm formation in the chronic colonization of CF patient lungs, as well as playing a role in infections of the cornea, burn wounds and chronic wounds. Our lab has previously characterized a secreted *P. aeruginosa* peptidase, PaAP, that is highly expressed in chronic CF isolates. This leucine aminopeptidase is not only secreted solubly, it also associates with bacterial outer membrane vesicles (OMVs), structures known for their contribution to virulence mechanisms in a variety of Gram-negative species and one of the major components of the biofilm matrix. With this in mind, we hypothesized that PaAP may play a role in *P. aeruginosa* biofilm formation. Using a lung epithelial cell/bacterial biofilm coculture model, we show that PaAP deletion in a clinical *P. aeruginosa* background leads to increased early biofilm formation. We additionally found that only native vesicle-bound PaAP, as opposed to its soluble forms, could reconstitute the original PaAP-mediated inhibition phenotype, and that the PaAP-containing vesicles could disperse preformed biofilm microcolonies of *Klebsiella pneumoniae*, another lung pathogen. These data provide the basis for future work into the mechanism behind PaAP-OMV mediated bacterial microcolony dispersal and the application of these findings to clinical anti-biofilm research.

## Introduction

The Gram-negative bacterium *Pseudomonas aeruginosa* is a prominent opportunistic pathogen capable of causing both acute and chronic disease in a variety of compromised hosts. *P. aeruginosa* is most well known as a leading cause of morbidity and mortality in cystic fibrosis (CF) patients^1^. In these individuals, the pathogen establishes chronic, biofilm-based infections that may persist for decades in the unique, mucous-rich environment of the CF lung. In addition to lung tissue, *P. aeruginosa* forms biofilms on a wide range of substrates relevant to human infection, including corneal and skin tissue, as well as soil and water reservoirs and hospital surfaces, which can contribute to infection initiation and spread^2^.

*P. aeruginosa* is considered a model organism for the study of biofilm formation, and many of the cellular and matrix components contributing to this mode of growth have been studied previously^3–5^. The process of forming biofilm communities has also been characterized, revealing distinct growth phases. First the bacteria settle and attach onto a suitable host tissue or abiotic surface, then they begin to form stationary microcolonies and secrete extrapolymeric substance (EPS) to form a dense matrix. This matrix contains a variety of polysaccharides, as well as proteins, lipids, OMVs, and eDNA^4,6^; however, the mechanisms by which many of these secreted bacterial components specifically affect microcolony growth and development remain unclear.

As with other extracellular pathogens, *P. aeruginosa* secretes many of its virulence determinants into the extracellular milieu to affect interactions with the host, including those in the biofilm matrix. Interestingly, many of the virulence factors secreted specifically by *P. aeruginosa* demonstrate proteolytic activities^7–9^. These enzymes, including elastase, protease IV, and alkaline protease, act on both bacterial and host proteins to directly impact host-pathogen interactions. For example, purified elastase has been found to cleave fibrin, laminin, various immunoglobulins, and components of the complement system^10^. This activity not only interferes with host defense mechanisms but may also impact host epithelial junctions, allowing the bacteria entrance to otherwise inaccessible tissues. These proteases can additionally aid in biofilm formation and antibiotic resistance^11^. Based on the importance of known virulence-associated, secreted proteases in *P. aeruginosa* biofilm formation, it follows that other proteases in the secretome, many of which have not been fully characterized, may also play a role in these processes.

With the goal of identifying factors that may contribute to chronic infections, our lab compared secretome protein expression profiles of clinical *P. aeruginosa* isolates from chronically infected CF patients to those of highly passaged laboratory and environmental strains. Several proteins were identified as being more highly expressed in all the clinical strains, indicating their potential involvement in virulence and chronic infection mechanisms. Among the proteins identified was aminopeptidase PA2939, known as PaAP (***<u>P</u>****seudomonas **<u>a</u>**eruginosa* **<u>a</u>**mino**<u>p</u>**eptidase)^12^.

Cahan et al. first described the proteolytic activity of PaAP in *P. aeruginosa* supernatants and classified it as a leucine aminopeptidase and a member of the M28 family of metalloproteases^9^. PaAP expression is regulated through quorum sensing (QS) mechanisms, and the protein is secreted via the *Pseudomonas* type II Xcp secretion pathway^13^. Initially, PaAP enters the extracellular environment as a 536 amino acid proenzyme. It is then processed by a variety of extracellular proteases at the C terminus to activate its enzymatic activity, and finally undergoes auto-processing at its N-terminus to create a 56 kDa, enzymatically active product^14^. It can be further processed extracellularly into several smaller, active products ranging from 55 kDa to 26 kDa. Outside of the cell, PaAP is found both soluble in the extracellular environment and as an abundant component of outer membrane vesicles (OMVs)^12^.

OMVs are formed from the envelope of Gram-negative bacteria and are small, discrete extracellular structures containing distinct membrane protein, lipid, and soluble periplasmic content^15^. They are known to interact with the external environment, including host tissues^16^. Vesicles produced by many Gram-negative pathogens, including *P. aeruginosa*, can elicit host immune responses, and many also serve as delivery mechanisms for bacterial virulence factors^16^. Of particular importance to our studies, OMVs have been implicated in the process of biofilm formation^6,15,17^. Schooling *et al.* demonstrated that OMVs are present in *P. aeruginosa* biofilm matrix and that bacteria in biofilms exhibit higher OMV production rates as compared to planktonic cells^18^. When these biofilm-derived vesicles were analyzed, PaAP was found to be one of the most abundant proteins ^19^.

Bacterial density, as communicated through quorum sensing, is known to be an important regulatory facet of biofilm development^20^. Not only is PaAP expression regulated by quorum sensing, the XCP type II secretion system that secretes PaAP is also QS-regulated^13^. In addition, when *P. aeruginosa* was cultured on host epithelial cells, PaAP expression was found to increase significantly^21^ suggesting that PaAP may play a role in biofilm formation on the host cells. Together, these data suggest PaAP’s potential role in fine-tuned pathogenesis processes, including biofilm formation and infection.

In this study, we report on the role of the secreted *P. aeruginosa* leucine aminopeptidase on biofilms formed on both abiotic and host cellular substrates. Using a coculture model in which single-species bacterial biofilms are formed on a confluent layer of host cells, we show that PaAP expression inhibits the formation of early biofilms on lung epithelial cells. Additionally, we characterize microcolony architecture and the vesicular components required for PaAP’s inhibitory activity against coculture biofilms. Finally, we investigate the broad impact of the PaAP-containing OMVs on biofilm development by non-self bacteria. These data provide novel insight into how the *P. aeruginosa* aminopeptidase can modulate clinically-relevant bacterial biofilms.

## Materials and Methods

### Bacterial strains and cell culture methods

*P. aeruginosa* strains used include the laboratory strain PAO1 (Pf1 phage-cured from our lab collection) and minimally passaged, non-mucoid cystic fibrosis clinical isolates CF2, CF3, and S470 as well as the previously-described isogenic mutants^12,22^. To generate fluorescent strains, we transformed bacteria with the dTomato expression plasmid, p67T1^23^. *Klebsiella pneumoniae* strain 43816 was obtained from ATCC. A549 human lung epithelia carcinoma cells (ATCC CCL-185) were grown in F-12K media containing 10% fetal bovine serum plus penicillin/streptomycin/fungizone. Unless specifically mentioned, all reagents were obtained from Sigma-Aldrich.

### Abiotic biofilm quantitation

Bacterial cultures were grown overnight in LB (Luria Broth (MILLER), Millipore-Sigma) (37°C, 200 rpm) with antibiotics if appropriate. The cultures were vortexed for 15 sec and diluted 1:1000 in M63 minimal media (supplemented with 1 mM magnesium sulfate, 0.2% glucose, and 0.5% casamino acids as outlined previously^24^) with no antibiotics. Aliquots of the diluted culture (100 mL/well) were incubated in five 96-well flat bottom polystyrene dishes (VWR) for 0, 2, 6, 10, or 24 h at 37°C with no shaking and evaluated using a static biofilm quantitation assay^24^. Briefly, at the incubation endpoint, the plate was washed out using distilled water (dH_2_0). After three washes, 150 mL of 1% aqueous Crystal Violet solution was added to each well and the samples were incubated for 10 min at room temperature. The stained wells were washed three times using dH_2_0, 200 mL of 30% aqueous acetic acid was added to solubilize the dye, and the OD_495_ was measured.

### Abiotic biofilm imaging

Glass coverslips were placed in 12-well polystyrene dishes (VWR) at an angle and sterilized with a UV light for 20 min. Bacterial cultures were grown (overnight, shaking, 37°C) in LB containing the appropriate antibiotics. Cultures were vortexed for 15 sec and diluted 1:1000 in M63 minimal media with no antibiotics. A portion (1 mL) was added to each well of the 12-well dish containing the coverslips, and the cultures were incubated for 5 or 24 h, without shaking at 37°C. The coverslips were removed from the cultures and briefly rinsed in dH_2_0. The coverslips were then mounted on glass slides and the biofilm structures examined using a Zeiss 780 inverted confocal microscope. Biomass was calculated using Comstat 2.1.

### qRT-PCR

Abiotic or coculture biofilms were grown for the indicated times as described. For coculture biofilms, supernatants were removed and 1 mL of Trizol was added to each sample to lyse both bacterial and A549 cells. For abiotic biofilms, 3 mL/well of cultures incubated in 6-well polystyrene dishes (VWR) were used to obtain sufficient biofilm material for analysis over the given time course. These samples were pipetted extensively to partially homogenize them and transferred to 15 mL conical tubes, in which the bacteria were pelleted. Trizol (1 mL, Invitrogen) was used to lyse each sample. A standard Trizol extraction was performed as described by the Invitrogen protocol, and RNA was extracted from the aqueous phase. The RNA was reprecipitated using 3 M sodium acetate for further purification and resuspended in dH_2_0. RNA samples were then DNase treated and reverse transcribed using the Applied Biosystems High-Capacity cDNA Reverse Transcription kit protocol. Applied Biosystems Power Sybr Green PCR Master Mix was used to prepare the samples, and they were analyzed on a StepOne Plus real-time PCR machine (Applied Biosystems). ΔC_T_ values were calculated relative to *proC* as a housekeeping control. Planktonic S470 ΔPaAP samples were used as a negative control, and relative expression was calculated against planktonic S470 WT samples, with samples prepared as for abiotic biofilm RNA from 5 mL overnight cultures in LB. Forward (F) and Reverse (R) primers used in this study: PaAP-F: GTGGTACGCAAGAAGACCGA, PaAP-R: ATCACCACGTTGTTCGGGTT, proC-F: CAGGCCGGGCAGTTGCTGTC, proC-R: GGTCAGGCGCGAGGCTGTCT^25^.

### Coculture assay

A549 cells were grown in MatTek glass bottom 3.5 mm tissue culture treated dishes for 10 days at 37°C in 5% CO_2_, with media changes every two days. Bacterial cultures were grown overnight in LB with the appropriate antibiotics, 1 mL of the culture was pelleted and resuspended in 1 mL sterile PBS (137 mM NaCl; 2.7 mM KCl; 10 mM Na_2_HPO_4_; 2 mM KH_2_PO_4_; pH 7.4). To fully resuspend the bacterial samples and reduce aggregation, each was passed through a 1 inch, 26-1/2G needle tip, 10 times. The OD_600_ was measured and conversion values were calculated to determine the number of cells/OD_600_ for each strain. For all following experiments, the OD_600_ was used to determine cell counts and multiplicity of infection (MOI). The bacterial samples were diluted in F-12K media (without antibiotic/antimycotic) and added to the A549-containing wells at MOI 30. The cocultures were incubated for 1h (37°C, 5% CO_2_) to allow the bacteria to attach to the cell surface. The cocultures were then washed three times with sterile PBS, and the media was replaced with F-12K containing 0.4% L-arginine. The cultures were incubated for an additional 4 h at 37°C and the media was replaced with microscopy-grade media (Sigma DMEM without Phenol Red, supplemented with 0.4% arginine). Images were taken on a Zeiss 780 inverted confocal microscope. Biomass calculations were completed using Comstat 2.1, and 3D colony models were created using the 3D viewer plugin for Image J. This protocol was adapted from previously published work^26^.

### Congo Red staining

To stain the biofilm extrapolymeric substance, a 100X Congo Red solution was made at 5 mg/mL in sterile PBS. The solution was sterile-filtered through a 0.22 mm PVDF syringe filter (VWR). Media on the biofilm cocultures to be stained was replaced with 1 mL of microscopy-grade media (Sigma DMEM without Phenol Red, supplemented with 0.4% arginine), and the Congo Red solution was added to a final concentration of 50 mg/mL. The samples were incubated at 37°C for 10 min, the staining media was removed, the samples were washed twice with PBS, and the media was replaced with fresh microscopy-grade DMEM (Sigma DMEM without Phenol Red, supplemented with 0.4% arginine). Microscopy images were then taken on a Zeiss 780 inverted confocal microscope. Biomass calculations were completed using Comstat 2.1.

### OMV isolation and supernatant fractionation

Bacterial cultures were grown overnight in LB at 37°C, diluted 1:100 into 1.5 L LB, and grown (37°C, shaking) to OD_600_ 0.9-1.1. Cells were pelleted (10,000×g for 15 min 4°C), and supernatants were collected and concentrated to 100 mL using tangential flow with a 10 kDa MWCO filter (Pall). The retentate was then filtered (0.22 mm, Pall) to generate cell-free supernatant (CFS). For detection of PaAP in CFS by immunoblotting, 1 mL of CFS was precipitated with 250 µL trichloroacetic acid (TCA), incubated at on ice for 1 hr, pelleted by centrifugation (12,500 × g), washed with 1 mL acetone, re-pelleted (12,500 × g) and resuspended in 0.75 M Tris-Cl, pH 8.8, and 1x SDS-PAGE sample buffer (1% β-mercaptoethanol, 0.004% bromophenol blue, 6% glycerol, 2% sodium dodecyl sulfate (SDS), 50 mM Tris-Cl, pH 6.8), prior to SDS-PAGE on BioRad 4-20% Tris-HCl gels. For anti-PaAP immunoblotting, gels were transferred to Amersham Hybond 0.45 µm polyvinyldifluoride (PVDF) membranes (GE Healthcare), blocked with 1% non-fat dry milk in TBS (50 mM Tris-Cl, pH 7.6; 150 mM NaCl), and then incubated (4°C, overnight) with anti-PaAP antibody^14^ diluted 1:1000 in TBS with 0.1% Tween (TBST). Blots were washed three times with TBST, incubated (room temperature, 1 h) with goat anti-rabbit Li-Cor secondary antibody (Odyssey) diluted 1:40,000 TBST + milk, washed, and analyzed using an Li-Cor Odyssey CLx imaging system with Li-Cor Image Studio software. ImageJ was used to quantify the anti-PaAP reactive bands.

To concentrate soluble proteins and OMVs from the CFS, ammonium sulfate was added to a final concentration of 90%, dithiothreitol added to 0.1 M, and the samples incubated overnight at 4°C with stirring. The precipitate was pelleted (15,000×g, 15 min, 4°C) and resuspended in 10 mL 20 mM HEPES, pH 8.0 (HEPES). The samples were then dialyzed in a ThermoFisher G2 cassette (10,000 MWCO) against HEPES, overnight at 4°C. The samples were further concentrated using Millipore 10,000 MWCO centrifugal filters, and the soluble proteins were separated from the OMVs using an iodixanol (Optiprep, Sigma) density gradient. Concentrated proteins and OMVs were adjusted to 50% Optiprep/HEPES in 2 mL, and applied to the bottom of the gradient tube, and subsequently layered with 2 mL of 40%, 2 mL of 35%, 4 mL of 30%, and 2 mL of 25% Optiprep/HEPES. The gradient was centrifuged overnight at 40,000×g, 4°C in an ultracentrifuge (SW 40 Ti swing bucket rotor, Beckman Coulter). Gradient fractions were removed from the top in 1 mL aliquots, and the lipid and protein content for each fraction were identified using FM4-64 and Bradford assays as described previously^27^. The OMV-containing fractions (typically 1-5) and soluble protein fractions (typically 7-12) were pooled and diluted with HEPES, then centrifuged at 40,000×g for 2 h to pellet OMVs and proteins and to remove Optiprep. Protein and OMV pellets were resuspended in HEPES, and sterile-filtered (0.4 mm PVDF, VWR).

### rPaAP purification

rPaAP expression, purification, and refolding were carried out as described previously^14^, and the refolded protein was concentrated using 10,000 MWCO Amicon filter units (Millipore-Sigma). The concentrated material was then dialyzed using the same method described above, applied to a S200 Sephadex ion exchange column, eluted with a gradient of 0-2 M NaCl, pH 7.6, at a flow rate of 0.02 mL/min, and 18- 2 mL fractions were collected. Fractions were assayed for leucine aminopeptidase activity as described^28^, and analyzed by SDS-PAGE and Ruby staining as described above. Fractions containing pure rPaAP were pooled and concentrated using Amicon 10,000 MWCO filter units to ∼0.5 mg/mL.

### Complementation and dispersion experiments

For biofilm complementation assays, CFS, OMV fractions, soluble fractions, and rPaAP samples were prepared as described above, total protein was measured by Bradford assay, and aminopeptidase activity was measured as described above. For standard complementation experiments, samples were added to cocultures with new media at 1 hpi. 50 ng total protein was used for CFS samples, and 25 ng total protein was used for OMVs, soluble fractions, and rPaAP samples. For co-addition experiments, 25 ng of PaAP^-^ OMVs were incubated with 25 ng of sPaAP or rPaAP for 10 min at room temperature prior to coculture treatment. In dispersion experiments, 25 ng of OMV samples were added to cocultures at 4.5 hpi. For OMV pretreatment of A549 cell layers, 25 ng of OMV samples were added to A549 cells for 15 min prior to inoculation. For experiments matched by PaAP activity, samples were standardized to the aminopeptidase activity found in 25 ng (by total protein) of PaAP^+^ OMVs.

### Statistics

For all experiments, statistics were completed using GraphPad Prism t-tests unless otherwise indicated. For microscopy images and imaging quantifications, representative results are shown.

## Results

### Leucine aminopeptidase expression does not affect biofilm formation on abiotic substrates

Based on previous reports describing the high expression of the *P. aeruginosa* aminopeptidase PaAP in both late-log phase and biofilm cultures, as well as the involvement of OMVs in the biofilm matrix, we wanted to determine whether the presence of PaAP was critical to biofilm formation. Several stages are observed during the process of *P. aeruginosa* biofilm growth and development. Bacteria initially attach to the substrate, then a variety of matrix components, including polysaccharides, proteins and eDNA, interconnect the microcolony structure, generating protection from environmental and immune stresses. As the microcolonies mature, bacterial cells disengage from the structure, to disperse and colonize in new environments^29^. When grown in liquid media, *P. aeruginosa* forms biofilm communities called pellicles at the air-liquid interface. As the bacteria attach to the culture container, this abiotic biofilm development can be quantitatively assessed using a Crystal Violet stain^24^.

To examine the effect of PaAP on *P. aeruginosa* pellicle growth, we compared a clinical isolate (S470 WT) that our lab previously found to exhibit high constitutive levels of PaAP expression, with the previously described isogenic knockout strain (S470 ΔPaAP) that contains a disruption of PA2939, the gene encoding PaAP^22^. Cultures of both strains were grown statically in 96-well dishes, and biofilm density was determined at various time points. We monitored growth up to 24 h when the biomass decreased, consistent with other static biofilm models of *P. aeruginosa*^24^. No significant differences were seen in the biofilms at any of the time points (**Fig 1A**) suggesting PaAP does not play a role in pellicle formation or stability.

**Figure 1:**
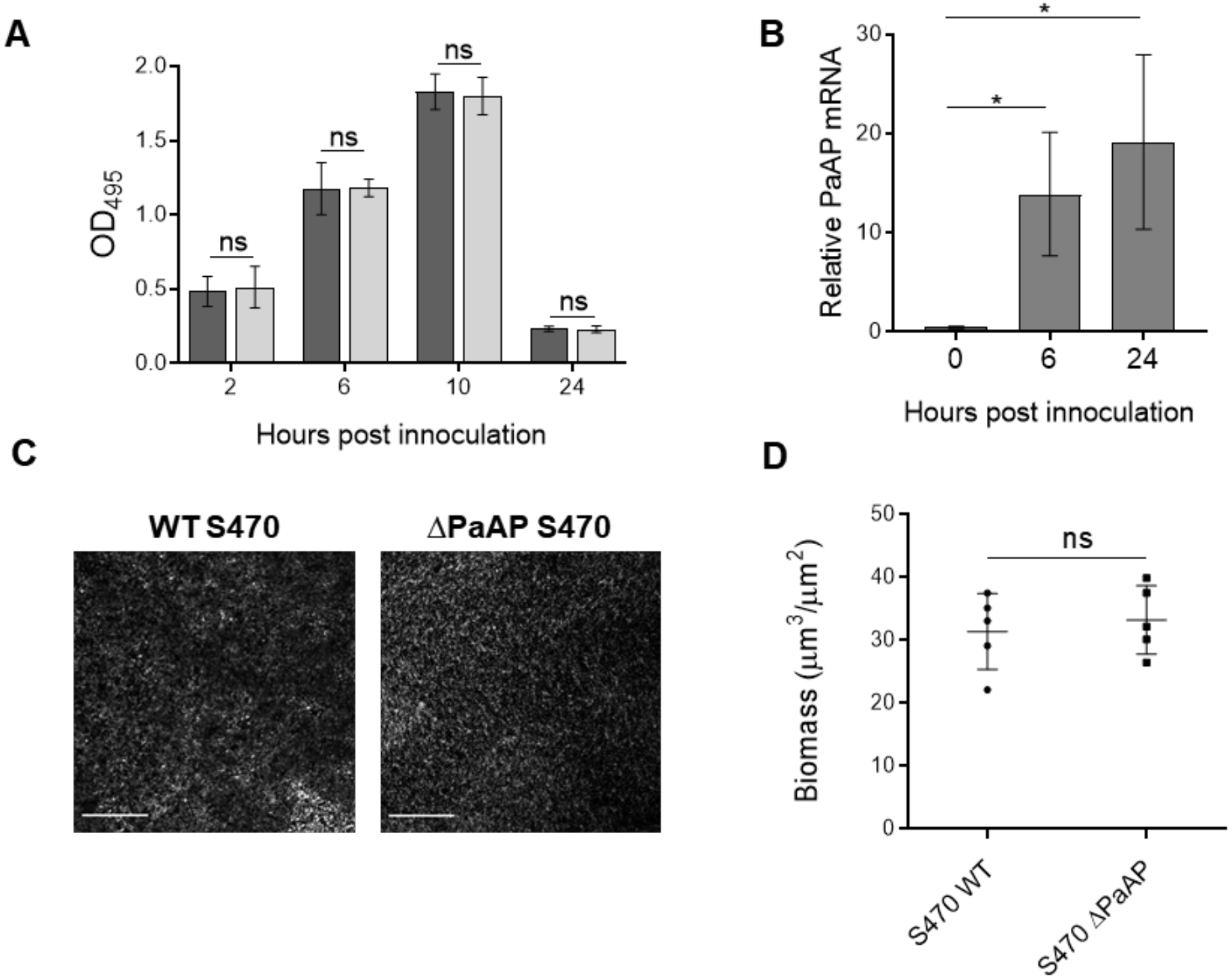
Deletion of the PaAP aminopeptidase had no effect on the structure or biomass of abiotic biofilms. **(A)** *P. aeruginosa* strains were grown in multi-well dishes for the indicated times and abiotic biofilms were quantified by Crystal Violet staining (OD_495_). (S470 WT: light gray; S470 ΔPaAP: dark gray) **(B)** Relative mRNA expression in S470 WT biofilms grown for the indicated times was calculated from qRT-PCR data using the ΔΔC_T_ method, with normalization to PaAP levels in planktonic S470 WT cultures. Pellicles formed by S470 WT or ΔPaAP (as indicated) on glass coverslips after 6 h of growth were imaged by confocal microscopy **(C)** and their biomass measured (D). For all experiments, n=3. *p<0.05; ns, not significant. Scale bars: 200 µm.

Because the Crystal Violet assay only measures cell density and not differences in biofilm structure, microcolony biomass and organization were also assessed by microscopy of pellicle biofilms using S470 WT and S470 ΔPaAP strains expressing the cytoplasmic fluorescent protein dTomato. These strains were grown in 12-well dishes containing glass coverslips inserted at an angle. Biofilms formed at the air-liquid interface attached to the coverslips and were examined using confocal laser scanning microscopy (CLSM). Qualitative examination of the microscopy images showed no differences between the strains with regards to the overall structure of the biofilms or organization of the microcolonies (**Fig 1C**). Total biovolumes for each image stack were calculated and the data confirmed that there were no significant differences between the wild type and PaAP-deficient strains (**Fig 1D**).

To confirm that the lack of a difference in biofilm phenotype observed in the above assays was not simply due to a lack of PaAP expression, PaAP RNA levels were measured in the static cultures of S470 WT and S470 ΔPaAP at 6 h and 24 h post infection (hpi) using qRT-PCR. As expected, the WT strains showed PaAP expression in the biofilms (**Fig 1B**), while the ΔPaAP S470 negative control did not (data not shown). Together, these data suggest that PaAP deletion does not impact the bacteria’s ability to develop a biofilm on abiotic substrates or at the air-liquid interface under our experimental conditions.

### *P. aeruginosa* early biofilm growth and microcolony organization on host epithelial cells is inhibited by PaAP expression

While PaAP did not affect the formation or architecture of *P. aeruginosa* biofilms on abiotic surfaces, we wanted to investigate whether PaAP affects biofilm formation on a live host cellular substrate as would be seen during *P. aeruginosa* acute or chronic respiratory infections. When *P. aeruginosa* was cultured on host epithelial cells, in addition to a variety of virulence-associated proteins, PaAP expression was found to increase significantly^27^, suggesting that PaAP may play a role in pathogenesis or biofilm formation on host cells. To assess differences in biotic biofilms, we adapted a previously developed biofilm coculture assay for use with A549 lung carcinoma cells^26^. When grown over the course of several days, these cells form fully confluent cell layers that partially model the polarized epithelial monolayers that the bacteria colonize during infection. Once grown to confluency they can be inoculated with bacteria and bacterial biofilm formation can be assessed by CLSM.

When confluent A549 cells were inoculated with the fluorescent S470 strain and incubated for up to 6 h, we noted significant damage to the A549 cells by 5 h, leading to breaks in the cell layer. This was consistent with the previously described toxicity of *P. aeruginosa* clinical strains to host cells^30^ and established time courses for static coculture biofilm models^26^. Once these cytotoxic bacteria infiltrated the host cell layer, the microcolony structure also broke down, as the bacteria were able to colonize the underlying tissue culture dish and migrate to the air-liquid interface. Thus, this cellular substrate could be used to investigate clinical *P. aeruginosa* strain biofilm development only through the early stages (up to 5 h post bacterial infection) of microcolony formation.

We quantitatively compared coculture biofilms of fluorescent S470 WT and S470 ΔPaAP strains at 5 hpi, and we observed significant differences between the two biotic biofilms. Surprisingly, the PaAP mutant strain formed biofilms with substantially greater cellular biomass compared to the WT strain (**Fig 2A,B**). Not only were these differences statistically significant, they were also clearly visible by qualitative examination of the fluorescent biofilms. These data provided the first evidence that PaAP modulates early biofilm growth under conditions mimicking host infection.

**Figure 2:**
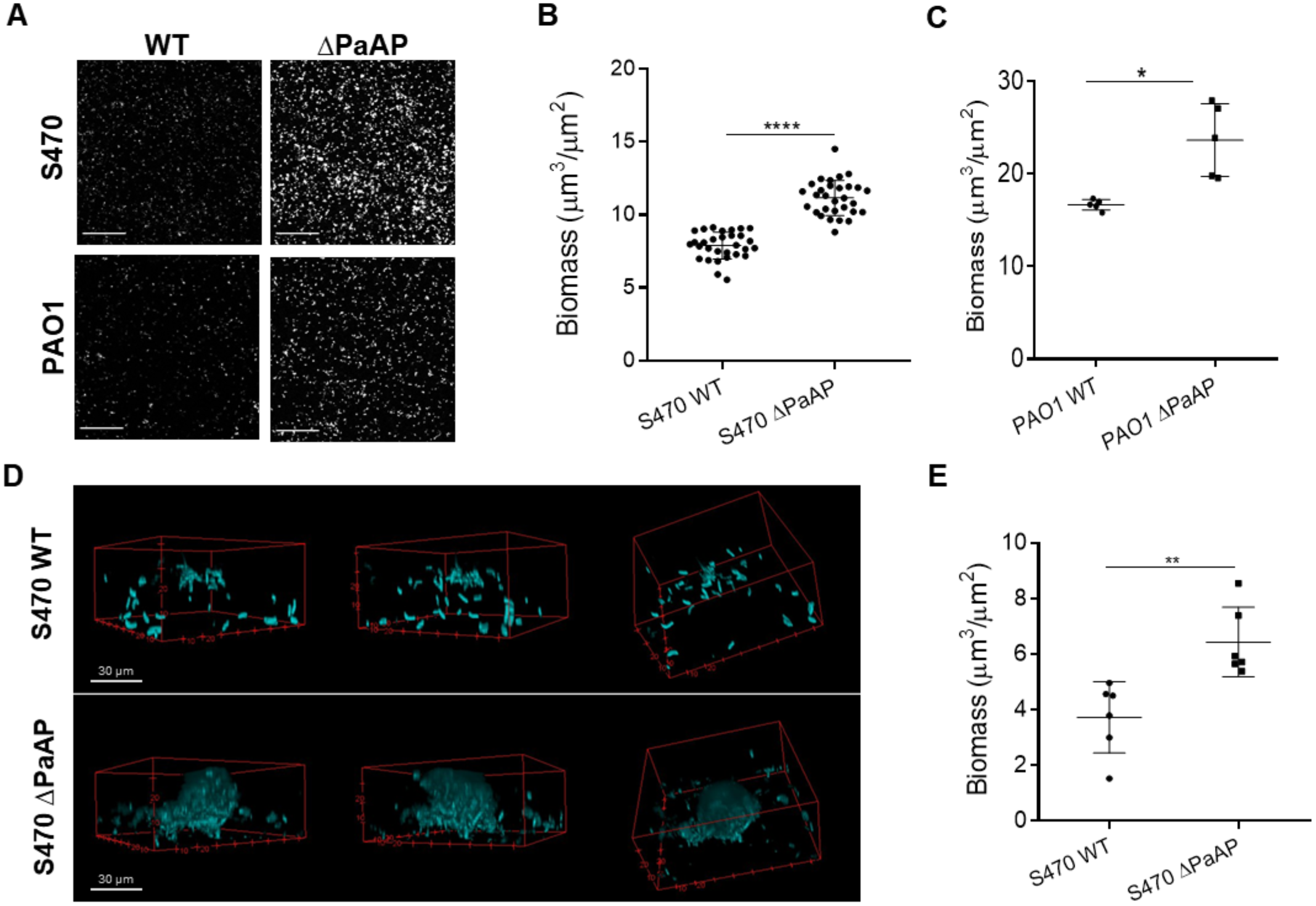
Deletion of the PaAP aminopeptidase increased the density, biomass, and organization of bacterial biofilms on host cells. Biofilms were grown on A549 confluent cell layers in glass bottom dishes. The protocol was adapted from a previous publication^27^. **(A)** S470 and PAO1 WT and ΔPaAP strains were inoculated onto cell layers, and images were taken at 10X magnification, 5 hpi and **(B, C)** the images were quantified. **(D)** For cocultures as described for A, images of microcolonies were taken using 100X magnification at 5 hpi and **(E)** quantified. For all experiments, n=3. *p<0.05, **p<0.01, ****p<.0001. Scale bars: 200 µm, unless otherwise specified.

To gain further insight into PaAP’s effect on biofilms, we examined S470 WT and ΔPaAP microcolony morphologies (**Fig 2D, E**). In addition to their greater quantitative biomass, S470 ΔPaAP colonies appeared to display greater cellular organization. Even at this early point in biofilm development, characteristic mushroom-shaped colony structures were observed for the PaAP mutant. The S470 WT microcolonies, by comparison, showed very little cellular organization and were packed less densely. Based on these data, we can conclude that the lack of PaAP not only increases cellular biomass, but also cellular organization at the microcolony level during early coculture biofilm development.

To confirm this PaAP-dependent phenotype in a different strain background, we compared the biofilm characteristics of PAO1, a laboratory *P. aeruginosa* strain commonly studied *in vitro*, and the isogenic PAO1 ΔPaAP strain that harbors a transposon insertion in PA2939. We note that our lab previously showed that OMVs isolated from PAO1 cultures show very low aminopeptidase levels compared to S470 and other clinical *P. aeruginosa* isolates^12^. Nevertheless, as with the clinical strain phenotype, PAO1 ΔPaAP formed significantly more biomass as compared to the WT counterpart (**Fig 2A, C**). Our observations indicate that PaAP is important to biofilm development, even in strains with low endogenous PaAP expression, and that its role may be relevant to a broad range of *P. aeruginosa* strains.

### Increased biofilm cellular density induced by PaAP deletion coincides with PaAP expression and increased EPS production

The results from the coculture assay in **Fig 2** revealed significant density and structural differences between the biofilms formed by WT and ΔPaAP *P. aeruginosa* strains based on bacterial fluorescence, and we were interested if PaAP also affected the overall extracellular, biofilm matrix. The *P. aeruginosa* matrix is composed of polysaccharides, proteins, DNA, and vesicles secreted by cells in the microcolony. Traditional imaging for biofilm matrices, such as eDNA staining methods, could not be used in our coculture system due to background staining of the underlying A549 cell layer. Instead, we adapted a Congo Red-based biofilm staining protocol to specifically label amyloid fibers and negatively charged extracellular polysaccharides in the *P. aeruginosa* matrix^31^. As shown in **Fig 3 A and B**, significantly higher levels of EPS staining were observed for S470 ΔPaAP cocultures than for the S470 WT counterpart. No staining was seen in the A549 cell-only controls. These data display the same trends observed for the S470 WT and ΔPaAP cellular biomass quantifications, supporting the discovery that PaAP expression prevents the formation of robust early biofilm structures.

**Figure 3:**
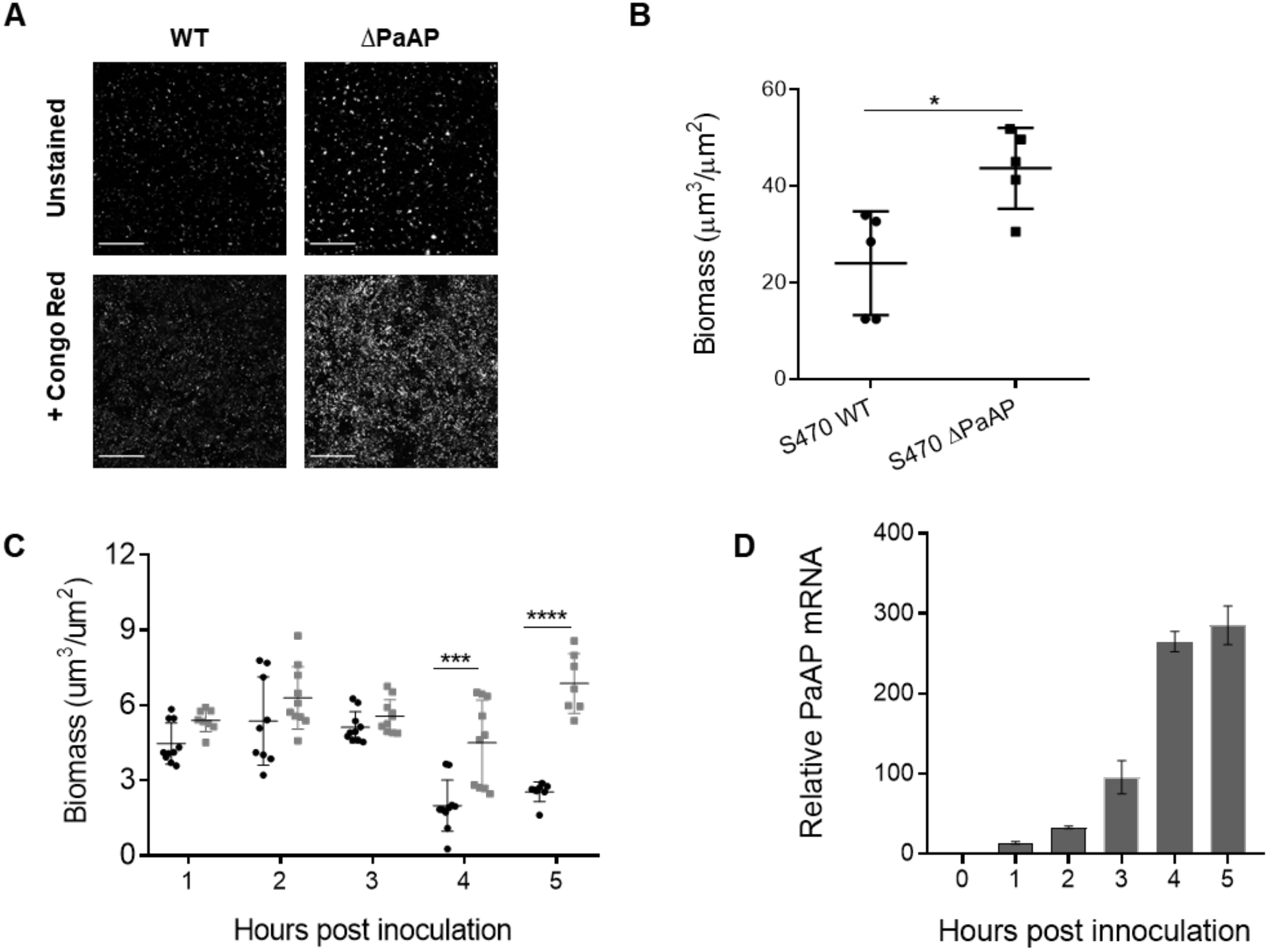
PaAP expression and the biofilm deletion phenotype develop coincidentally during the observed time course. Bacterial biofilms were cocultured on A549 cells for 5 h, stained with Congo Red, imaged by confocal microscopy **(A)** and total biomass was calculated **(B)**. **(C)** Bacterial biofilms cocultured on A549 cells were imaged by confocal microscopy at indicated time points. (S470 WT: dark gray, S470 ΔPaAP: light gray). **(D)** For cocultured biofilms of S470 WT, PaAP mRNA expression was measured at the indicated time points using qRT-PCR and calculated using the ΔΔC_T_ method, with normalization to PaAP levels in planktonic S470 WT cultures. For all experiments, n=3. *p<0.05, ***p<0.001, ****p<0.0001. Scale bars: 200 µm.

There are several mechanisms through which PaAP may be able to affect the formation of early biofilms, including preventing efficient attachment of bacteria to the host cellular substrate and affecting colony growth post-attachment. To detail the time course of the PaAP- dependent phenotype, we imaged the cocultures at hourly intervals post infection. Directly following the wash step at 1 hpi and up to 3 hpi, we observed a similar level of biomass from the S470 WT and ΔPaAP strains. A significant phenotype did not develop until 4 hpi (**Fig 3C**), suggesting that the aminopeptidase functions to limit microcolony growth and development, rather than the initial steps of colonization and attachment.

Based on this timeline, it is expected that the observed phenotype developed in parallel with the level of aminopeptidase expression. Indeed, previous reports found PaAP to be regulated by the *las* quorum sensing system^13,32^, and thus PaAP expression would be expected to increase as bacterial density increases in our experimental system. Using qRT-PCR, we examined the expression of PaAP in S470 WT biofilms over time under the coculture assay conditions. Consistent with quorum sensing density-dependent expression, our results show expression of PaAP mRNA at 1 hpi, increasing steadily over the next several hours (**Fig 3D**). Based on these data, it appears that the biofilm inhibition phenotype may depend on either a threshold level of aminopeptidase or a specific microcolony developmental phase.

### Addition of PaAP-containing outer membrane vesicles, but not purified aminopeptidase, inhibits biofilm development by the PaAP deletion mutant

Based on its tightly-controlled, quorum sensing-regulated expression and secretion pathways, PA2939 complementation using plasmid-based expression has proven to be challenging and unfeasible; therefore, we performed biochemical complementation experiments. Complementation was first tested using cell-free supernatants from mid-log phase cultures of S470 WT and S470 ΔPaAP. We confirmed the presence and activity of PaAP in the S470 WT supernatant and its absence in the S470 ΔPaAP supernatant by Western blot (**Fig 4A**) and aminopeptidase activity assays (data not shown). Equivalent amounts (by total protein) of the sterile supernatants, from S470 WT and ΔPaAP cultures were added to the cocultures of S470 ΔPaAP and A549 cells after the 1 hpi wash step, and incubated until the 5 hpi final imaging time point. Addition of the PaAP-containing S470 WT supernatant significantly decreased the bacterial biovolume in the coculture (**Fig 4B**), while the S470 ΔPaAP supernatant had only minor qualitative effects on colony size and shape. These results provide strong evidence that secreted aminopeptidase is responsible for PaAP-dependent biofilm inhibition.

**Figure 4:**
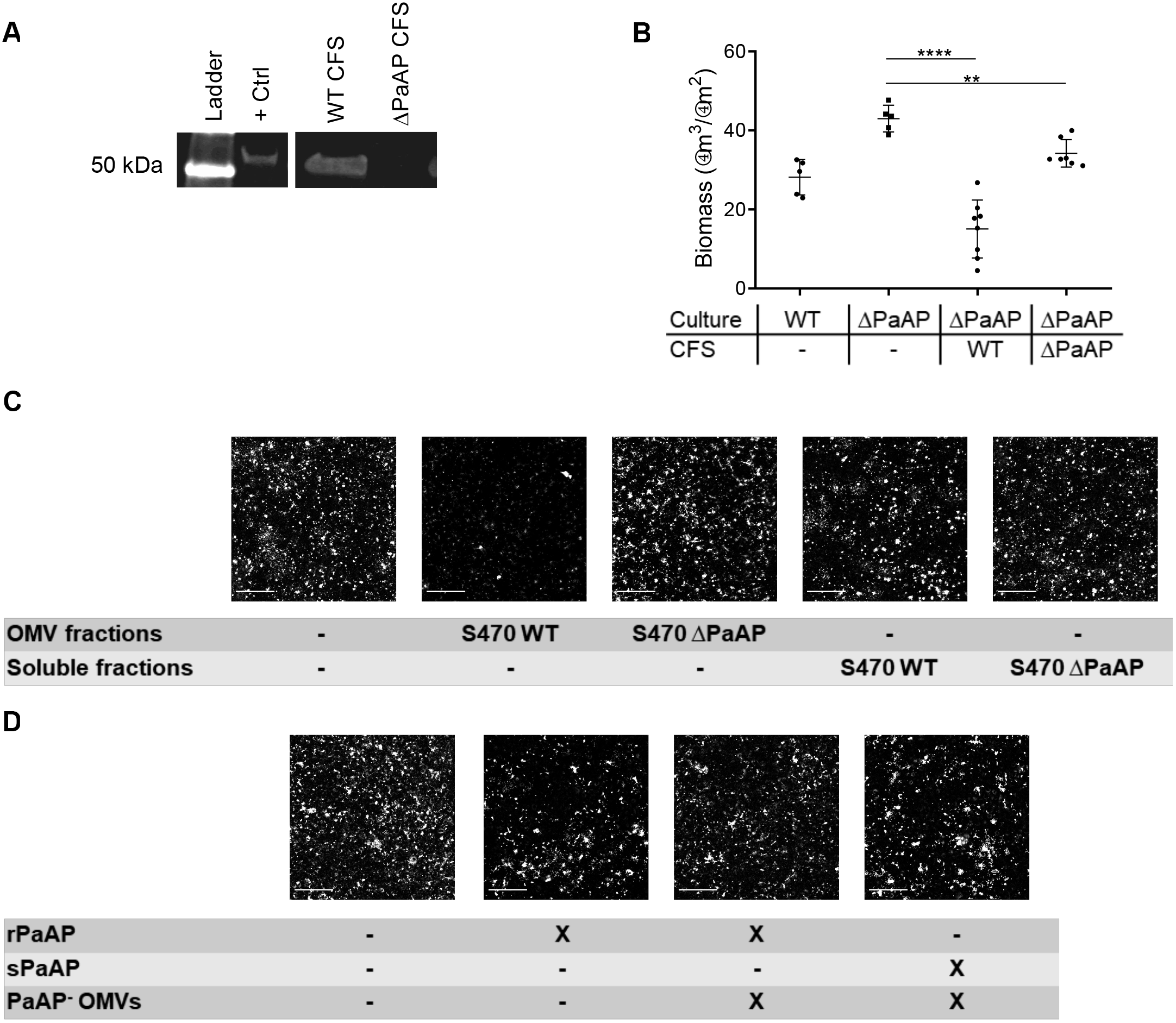
Addition of PaAP^+^ OMVs inhibited formation of *P. aeruginosa* coculture biofilms. **(A)** Cell free supernatants from S470 WT (WT CFS) and S470 ΔPaAP (ΔPaAP CFS) were TCA-precipitated, and aminopeptidase was detected by immunoblotting after separation of samples by SDS-PAGE. A lane with rPaAP was run as a positive control (+Ctrl), and the migration of the 50 kDa molecular weight standard is shown. **(B)** S470 ΔPaAP (ΔPaAP) biofilms cocultured with A549 cells were treated with CFS S470 WT or ΔPaAP at 1 hpi and the biomass of these samples and the S470 WT untreated control were quantified. **(C)** S470 ΔPaAP biofilms cocultured on A549 cells were treated at 1 hpi with pooled Optiprep fractions containing either OMVs or soluble products from S470 WT or S470 ΔPaAP CFS as indicated. (D) S470 ΔPaAP cocultures were treated at 1 hpi with combinations of rPaAP, PaAP^-^ soluble fractions, and/or PaAP^-^ OMV fractions from S470 ΔPaAP CFS as indicated. For all experiments, n=3. **p<0.01, ****p<0.0001. Scale bars: 200 µm.

The addition of WT cell-free supernatant to the ΔPaAP coculture mimics the effect of endogenously expressed and secreted PaAP; however, WT *P. aeruginosa* supernatants contain both secreted soluble and OMV-bound forms of the aminopeptidase^12^. To identify which form of PaAP in the S470 WT supernatant was responsible for the observed decrease in S470 ΔPaAP biofilm development, soluble proteins and OMV-bound factors were separated using iodixanol density gradients. Light density fractions from these gradients are enriched in OMVs, whereas more dense fractions are enriched in soluble proteins^28^. Also consistent with our previous reports, aminopeptidase activity was detected in both the light and heavy fractions, representing the OMV-bound (PaAP^+^ OMVs) and soluble forms of PaAP (sPaAP), respectively. The ability of equivalent amounts (by total protein) of the light and heavy density fractions to inhibit S470 ΔPaAP biofilms was assessed. Intriguingly, while the OMV-containing S470 WT fractions were able to significantly inhibit biofilm development by S470 ΔPaAP, the heavier, soluble aminopeptidase-containing fractions had no noticeable effect on biomass (**Fig 4C**). Experiments using the different density fractions matched to aminopeptidase activity levels found in PaAP^+^ OMVs yielded similar results (data not shown), confirming that, regardless of enzymatic activity, the soluble material was unable to inhibit biofilm growth. These data suggest that PaAP- containing OMVs, but not soluble secreted aminopeptidase, inhibit early stages of biofilm development.

To further corroborate our finding that soluble forms of PaAP cannot prevent microcolony development, we purified and refolded recombinant *P. aeruginosa* aminopeptidase (rPaAP) expressed in *E. coli*^14^. The specific activity of this preparation was similar to, though slightly lower than, that of the native soluble enzyme purified from culture supernatants. Equivalent amounts (by activity) of rPaAP and PaAP+ OMVs were added to the S470 ΔPaAP cocultures to test their ability to modulate biofilm formation. Consistent with the results using sPaAP, purified rPaAP was also incapable of disrupting the formation of robust biofilms by the ΔPaAP strain (**Fig 4D**).

We then hypothesized that addition of rPaAP or sPaAP may reconstitute the biofilm inhibitory activity of the OMVs isolated from ΔPaAP cultures (PaAP^-^ OMVs). These rPaAP/PaAP^-^ OMV and sPaAP/PaAP^-^ OMV preparations had demonstrable aminopeptidase activity (data not shown), but the co-additions failed to inhibit biotic biofilm development by the S470 ΔPaAP strain (**Fig 4D**). It may be that interactions between the aminopeptidase and the PaAP^-^ OMVs were not natively reconstituted in our co-incubated preparations, as characteristics of rPaAP and sPaAP possibly prevent their interactions with OMVs. Nevertheless, these data suggest that native secretion and/or association with OMVs is critical for PaAP-dependent biofilm inhibition.

To explore which components of the vesicles were critical to PaAP^+^ OMV-mediated biofilm inhibition, OMVs were treated to disrupt the native vesicle structure. These results are summarized in **Table 1**. Treatment with Triton X-100 is expected to disrupt the lipid membrane structure and effectively liberate OMV cargo. Unexpectedly, detergent treatment significantly increased overall biofilm formation in both the S470 WT and ΔPaAP OMV-treated samples. It is unclear what caused this, as both soluble and insoluble factors could be liberated by detergent; however, it prohibited us from assessing the effect of PaAP in a “burst vesicle” model.

**Table 1:** OMV-bound PaAP enzymatic activity and biofilm inhibition phenotypes are resistant to most physical and protease treatments

| Treatment<sup>1</sup> | OMVs | AP Activity<sup>2</sup> | Biomass<sup>3</sup> | Interpretation |
| --- | --- | --- | --- | --- |
| None | None | - | = | Untreated biofilm and untreated OMV controls |
|  | S470 WT | 100.0 (±4.4) | - - |  |
|  | S470 ΔPaAP | - | = |  |
| Triton X-100 | S470 WT | 296.7 (±3.1) | + | OMV disruption by detergent increases biomass in a PaAP-independent manner, preventing assessment of Triton X-100 treatment on complementation |
|  | S470 ΔPaAP | - | + |  |
| Proteinase K | S470 WT | 113.4 (±2.5) | - - | Proteinase K cannot access PaAP to fully degrade the enzyme, but may enhance PaAP activity through enzymatic activation |
|  | S470 ΔPaAP | - | = |  |
| Boil (95°C) | S470 WT | 36.3 (±4.5) | - - | Degradation of PaAP and the vesicle structure occurs, but aminopeptidase and dispersal activities remain, indicating PaAP is somewhat protected |
|  | S470 ΔPaAP | - | = |  |
| Boil (95°C), Proteinase K | S470 WT | 8.6 (±3.9) | - | Boiling allows Proteinase K access to vesicle-associated PaAP, but limited aminopeptidase and dispersal activities remain, showing residual PaAP protection |
|  | S470 ΔPaAP | - | = |  |
| Boil (95°C), Triton X-100, Proteinase K | S470 WT | 1.3 (±0.6) | = | Boiling and detergent treatments release PaAP from protective elements, granting Proteinase K full access to degrade remaining PaAP and eliminate anti-biofilm activity |
|  | S470 ΔPaAP | - | = |  |

<sup>1</sup>S470 WT and ΔPaAP OMVs were treated, as indicated, with combinations of 0.1% Triton X-100, 100 µg/mL Proteinase K, or boiling for 15 min at 95°C. Proteinase K and Triton X-100 treated samples were incubated at for 1 h at 37°C. Combinatory treatments were done in the indicated order.
<sup>2</sup>Aminopeptidase activity levels (±SD) were measured post-treatment, and values were normalized to S470 WT OMV aminopeptidase activity.
<sup>3</sup>OMV samples were added to S470 ΔPaAP biofilm cocultures 1 hpi and biomass was quantified 5 hpi. Biomass was calculated relative to untreated S470 ΔPaAP cocultures (=: no significant increase; +: significant increase; -: significant decrease). The number of +/- quantifies the relative biomass differences (+/-: 5-25%; +/- -: 26-50%; +++/- -: >50%).

To investigate if externally-localized protein is necessary for PaAP^+^ OMV-mediated biofilm inhibition, we treated samples with a membrane impermeable enzyme, Proteinase K. We found that Proteinase K addition increased the leucine aminopeptidase activity of the vesicle samples. We note that this was not due to an off-target effect of Proteinase K on the aminopeptidase assay substrate, as an increase in cleaved substrate was not seen in any control samples, and sPaAP samples were fully degraded as expected (data not shown). Based on the mechanism of PaAP’s extracellular processing and activation by other proteases^14^, it follows that increased aminopeptidase activity may be caused by partial Proteinase K cleavage of PaAP, leading to PaAP activation rather than expected degradation. This suggests that Proteinase K has partial access to the aminopeptidase to cause activation, leading us to conclude that it is not fully protected by the vesicle membrane. However, PaAP is protected from full degradation by its association with the vesicle, and anti-biofilm activity was not affected by the treatment.

In an attempt to eliminate full aminopeptidase and anti-biofilm activity from PaAP^+^ OMVs, the vesicles were boiled at 95°C for 15 minutes. This treatment is expected to degrade all vesicular structures, membranes, and proteins; however, nearly one third of aminopeptidase activity and some anti-biofilm activity remained. A combination of boiling and Proteinase K further decreased aminopeptidase and anti-biofilm activity in the samples, but full degradation was not observed until the OMVs were treated with boiling, Triton X-100, and Proteinase K, which effectively liberated the aminopeptidase from any protective elements, allowing digestion. The highly resistant nature of the aminopeptidase-vesicle association strengthens our conclusion that PaAP anti-biofilm activity is innately linked to native packaging into outer membrane vesicles. Additionally, the anti-biofilm activity of these particles directly paralleled their aminopeptidase activity, providing further evidence that PaAP mediates OMV anti-biofilm activity.

### PaAP^+^ OMVs disperse early *P. aeruginosa* and *K. pneumoniae* biofilms

Having determined that PaAP-containing OMVs are able to inhibit coculture biofilm formation, we were curious at what point in microcolony development these particles act. To investigate this, OMVs were added to cocultures either before inoculation (pre-treatment), at 1 hpi, or 0.5 h before the final imaging time point. **Table 2** summarizes the findings of these experiments. The results using A549 cell layers pretreated with OMVs prior to bacterial addition allowed us to determine whether the vesicles interact with the host cells to prevent efficient bacterial attachment to the host surface. Differences between the PaAP^+^ and PaAP^-^ OMV pre-treated cocultures did not develop until late time points, providing evidence that the PaAP^+^ OMVs did not influence the ability of bacteria to attach to the host cell surface. Consistent with an effect on microcolony development rather than attachment, PaAP^+^ OMV treatments 1 h after S470 ΔPaAP bacterial inoculation resulted in substantially reduced microcolony structures at the 3 h and 5 h post-infection time points.

**Table 2:** PaAP^+^ OMVs inhibit microcolony development and disperse pre-formed microcolonies

| OMV Addition<sup>1</sup> | Imaging Time<sup>2</sup> | OMVs | Biomass<sup>3</sup> | Interpretation |
| --- | --- | --- | --- | --- |
| Bacterial Attachment |  |  |  |  |
| 0 hrs<br>(pre-treatment) | 1 hr | None (control) | = | OMV pretreatment of the A549 cells does not affect initial bacterial cell attachment to the host surface. |
|  |  | S470 WT | = |  |
|  |  | S470 ΔPaAP | = | This suggests PaAP's mechanism does not affect bacteria/host cell initial adhesion |
| Microcolony Inhibition |  |  |  |  |
| 0 hrs<br>(pre-treatment) | 5 hrs | None (control) | = | Addition of PaAP-containing OMVs at any point early in the coculture experiment results in lower bacterial biomass. |
|  |  | S470 WT | - - |  |
|  |  | S470 ΔPaAP | = | This effect is more pronounced at later time points, suggesting PaAP-mediated biofilm inhibition acts on a bacterial pathway important to microcolony growth and formation, rather than initial attachment. |
| 1 hr | 3 hrs | None (control) | = |  |
|  |  | S470 WT | - |  |
|  |  | S470 ΔPaAP | = |  |
| 1 hr | 5 hrs | None (control) | = |  |
|  |  | S470 WT | - - |  |
|  |  | S470 ΔPaAP | = |  |
| Microcolony Dispersion |  |  |  |  |
| 4.5 hrs | 5 hrs | None (control) | = | PaAP bound OMVs can disperse established microcolonies, further suggesting that the aminopeptidase affects microcolony growth and dispersal mechanisms. |
|  |  | S470 WT | - - - |  |
|  |  | S470 ΔPaAP | = |  |

<sup>1</sup>S470 ΔPaAP cocultures were treated with the indicated OMVs at the indicated times. For pre-treatment, A549 cells were incubated with vesicles for 0.5 h before bacterial inoculation.
<sup>2</sup>Samples were imaged at the indicated times and assessed for biomass.
<sup>3</sup>Biomass is represented relative to the untreated S470 ΔPaAP controls for each experiment (=: no significant difference; +: significant increase; -: significant decrease). The number of +/- quantifies the relative biomass differences (+/-: 5-25%; +/- -: 25-50%; +++/- -: >50%).

To test directly whether PaAP-containing vesicles can act to disrupt preformed microcolony structures, OMVs were added to the A549/S470 ΔPaAP cocultures at 4.5 h post-infection, 30 minutes before the final imaging time point. As seen in **Fig 5A**, the PaAP- containing OMVs not only complemented the knockout phenotype, but the cellular biomass was virtually non-existent in these samples. To demonstrate that this phenotype was not specific to S470-derived vesicles, it was recapitulated with PAO1-derived OMV treatment of A549/S470 ΔPaAP cocultures. This treatment also led to biofilm dispersion; however, consistent with lower aminopeptidase expression by the PAO1 strain, PAO1 WT OMVs dispersed S470 ΔPaAP biofilms to a much lesser extent (**Fig 5A**). Taken together, these experiments provide evidence that OMV-associated PaAP is able to disrupt microcolonies already formed on biotic substrates in addition to inhibiting their formation.

**Figure 5:**
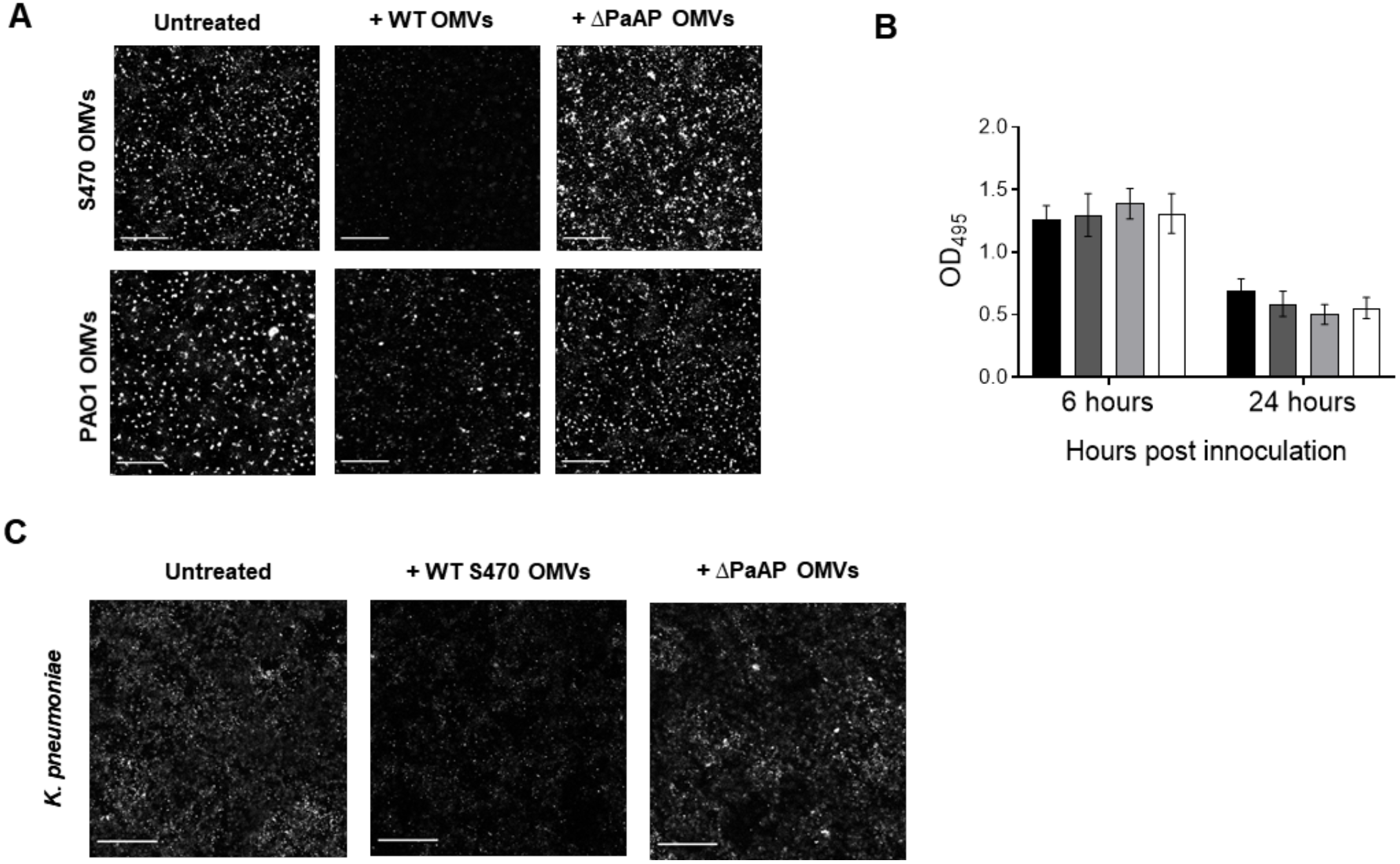
PaAP-bound OMVs disperse pre-formed biofilms from both pseudomonads and non-pseudomonads. **(A)** Top row: OMVs from S470 WT or ΔPaAP cultures were added to S470 ΔPaAP biofilm cocultures at 4.5 hpi, 30 minutes prior to imaging. Bottom row: OMVs from PAO1 WT or ΔPaAP cultures were added to S470 ΔPaAP biofilm cocultures at 4.5 hpi, 30 minutes prior to imaging. **(B)** S470 WT or ΔPaAP OMVs were added to *P. aeruginosa* cultures in polystyrene dishes 4.5 hpi. CV staining was used to assess biofilm formation. (S470 WT untreated: black bars; S470 ΔPaAP untreated: dark gray bars; S470 ΔPaAP + WT OMVs: light gray bars; S470 ΔPaAP + ΔPaAP OMVs: white bars). **(C)** *K. pneumoniae* cocultures were treated with S470 WT or ΔPaAP vesicles at 4.5 hpi and biofilm formation was assessed by Congo Red staining at 5 hpi. For all experiments, n=3. p-values were not significant. Scale bars: 200 µm.

To further characterize the biofilm modulatory capabilities of PaAP-containing OMVs, we also tested their ability to inhibit or disperse biofilms grown on abiotic substrates in multi-well dishes. Equal amounts (by protein) of S470 WT and ΔPaAP OMV preparations were added to S470 ΔPaAP strains either during inoculation of the static cultures or 0.5 h prior to staining. At 2, 6, and 24 hpi, biofilm formation was assessed using a Crystal Violet stain. As shown in **Fig 5B**, no significant difference was seen between the untreated controls and the vesicle-treated biofilms in either the inhibition or dispersion Crystal Violet experiments. It should be noted that these results were not surprising, considering the lack of phenotypic differences between the wild type and the PaAP mutant using the abiotic biofilm assay.

During infections, such as occur in the lungs of CF patients, biofilms are often found as polymicrobial communities. As such, components secreted by one species can impact the microcolonies of other strains or species in nearby environments. Thus far, we have shown that PaAP^+^ OMVs prevent and disperse biofilms of *P. aeruginosa*. To test if they could also exhibit cross-species effects, A549 cell layers were inoculated with *K. pneumoniae*, a bacterium which is often found with *P. aeruginosa* in cases of ventilator-associated pneumonia^33^. S470 WT or ΔPaAP OMVs were added at 4.5 h post-infection, and the cocultures were stained at 5 h with Congo Red to examine both cellular and matrix biomass. As seen in **Fig 5C**, S470 WT vesicles significantly reduced *K. pneumoniae* total biomass, while ΔPaAP OMVs had very little effect. These data suggest that the OMV-associated *P. aeruginosa* aminopeptidase not only acts to disrupt a process that is critical to the development of coculture biofilms by *P. aeruginosa*, but also that it has the ability to impact the growth and development of another clinically relevant pathogen.

## Discussion

We set out to investigate the role of the PaAP leucine aminopeptidase in biofilm development, as PaAP is one of the most abundant proteins in the *P. aeruginosa* biofilm extracellular matrix and one of the major protein components of OMVs from clinical *P. aeruginosa* isolates^12,19^. In this study, we have shown that deletion of the PaAP-encoding gene did not affect the ability of clinical or laboratory *P. aeruginosa* strains to form biofilms on a polystyrene surface. However, when the bacteria were grown on a host cellular substrate, the absence of PaAP significantly increased cellular and matrix biomass, as well as microcolony density and organization. We discovered that the addition of the OMV-containing fractions of wild type bacterial culture supernatants could reduce biofilm formation by the mutant bacteria. Further, we showed that purified OMVs from PaAP-expressing strains not only were able to inhibit the formation of early biofilms of PaAP-deficient *P. aeruginosa*, but they could also disperse biofilms that had reached early levels of maturity. Their inhibitory activity was highly resistant to degradation, and neither high density fractions containing native soluble active aminopeptidase, nor purified active recombinant PaAP, nor a mixture of aminopeptidase with PaAP^-^ OMVs exhibited these inhibition and dispersion properties. These results suggest that not only PaAP, but also the native vesicle context of PaAP, is critical to exert this behavior. Finally, we provided evidence that PaAP-containing OMVs could disperse coculture biofilms of heterologous bacterial species, specifically, pre-formed *K. pneumoniae* biofilm microcolonies. Our work supports a hypothetical mechanism in which the aminopeptidase and the native vesicle structure function synergistically to target bacterial microcolonies forming on host cells, aiding in biofilm inhibition and dispersion.

A quite distinct role for PaAP in *P. aeruginosa* biofilm development has been reported recently. Zhao *et al.* published data showing that deletion of PaAP resulted in abiotic biofilm phenotypes: an increase in early PAO1 abiotic biofilm growth (up to 6 h post inoculation) and a reduction at later time points (36-48 h)^34^. Consideration of their experimental design shows why our results, which did not include an abiotic phenotype, and the mechanistic roles for PaAP revealed in our two studies differ substantially. First, the bacteria in our experimental assays had access to more complex carbon sources than the cultures used by Zhao *et al.*, and second, our assays were performed at 37°C instead of 30°C. Zhao *et al.* used Jensen’s media (a bacterial growth media designed to increase *P. aeruginosa* elastase expression^35^). For the data presented in this report, biofilms were grown in M63 minimal media supplemented with casamino acids and glucose; however, we note that we also compared abiotic biofilms of WT and mutant strains grown in LB, LB/glycerol, and A549 cell culture media, with no observed differences in phenotype (data not shown). Combined with the difference in growth temperature, these nutritional differences would substantially impact the metabolic profile of the biofilm bacteria and reveal the distinct role of PaAP under Zhao’s experimental conditions. In those assays, PaAP’s activity is likely required for bacterial growth due to its ability to generate amino acid-based carbon sources. Indeed, they discovered that during late growth phases, the absence of PaAP led to death and lysis of *P. aeruginosa*. Consequent to lysis, the cells released a periplasmic enzyme, PslG, that enabled biofilm matrix digestion, a mechanism that is likely used in the environment to liberate bacteria so that they can reach a more nutrient-rich niche. By contrast, in our assay conditions the growth and death phases of WT and ΔPaAP cultures were indistinguishable (**Fig 1A**). In sum, the mechanistic role for PaAP in *P. aeruginosa* biofilm modulation described in these two studies appear to be substantially different: in one, PaAP activity prevents starvation during nutrient deprivation and prevents the otherwise death-dependent release of an abiotic biofilm-degrading intracellular enzyme, and in the other, a form of the secreted aminopeptidase has a biotic biofilm inhibiting and degrading activity.

Since PaAP was known to exist in two enzymatically active secreted states, OMV-bound and soluble, we were curious whether one or both were active in dispersing biofilms. The light density OMV-containing fraction, but not the heavier sPaAP-containing fraction of the cell free supernatant of WT *P. aeruginosa* was found to harbor the early biofilm inhibitory activity. In addition, the combination of rPaAP to ΔPaAP OMVs failed to inhibit or disperse S470 ΔPaAP biofilms. These data suggested that subtleties of native vesicle packaging were critical, potentially by impacting PaAP orientation or association within the vesicle and vesicle-mediated delivery to the biofilm matrix. While future work will be required to determine PaAP’s localization with regard to the vesicle structure, preliminary data from our lab, along with the fact that PaAP is known to be secreted through the OM, suggest that during OMV biogenesis the aminopeptidase localizes to the surface of OMVs^22^.

The inability of PaAP^+^ OMVs to affect biofilms grown under our abiotic experimental conditions while dispersing partially maturing coculture biofilms of both pseudomonads and non-pseudomonads generated some further insight into the dispersal mechanism. The data do not favor the possibility that the differences in biofilms generated by the wild type and mutant bacteria are dependent on differences in host-derived biofilm factors secreted differentlially in response to the two bacterial strains. First, there is no significant difference in cytokine expression responses elicited for incubations with the S470 WT and ΔPaAP bacteria (data not shown), and second, the WT OMVs are active towards pre-initiated ΔPaAP mutant bacterial biofilms. Further, as we also observed activity against non-pseudomonas biofilms, it appears that the factor targeted by the PaAP^+^ OMVs is a generally conserved feature of bacterial biotic biofilms. Together, these data lead us to favor a model whereby PaAP^+^ OMV-mediated biofilm dispersal relies on targeting a factor (or factors) governing biofilm integrity that is either conserved and host cell-derived or conserved and host cell-elicited, but bacterially-derived, which would be consistent with the activity of many anti-biofilm enzymes against the biofilm matrix^36^.

Our data addressing the timing of PaAP^+^ OMV-mediated biofilm dispersal support the following model, which outlines OMV interactions during the microcolony development stages observed in our coculture experiments (**Fig 6**). The structural differences between these phases directly impact the mechanism by which PaAP^+^ OMVs can access and disperse biofilms. Pretreatment of the A549 host cell layer did not affect initial bacterial attachment, suggesting that direct PaAP^+^ OMV-host cell interactions likely are not responsible for the observed dispersal phenotype. Rather, PaAP^+^ OMVs demonstrated anti-biofilm activity at late stages of coculture infection, at which point they likely have decreased access to the host cells and act to disperse a developing matrix structure and multi-cellular architecture. PaAP^+^ OMV production induced by quorum sensing inside the biofilm by WT *P. aeruginosa* would exhibit biofilm inhibiting activity at this point. Further, based on the ability of exogenously added PaAP^+^ OMVs to disperse pre-formed *P. aeruginosa* and *K. pneumoniae* microcolonies, we note that released PaAP^+^ vesicles can target and dissociate developing biotic microcolonies that they encounter in the environment. It is widely appreciated that OMVs can serve as vehicles to deliver bacterial toxins and effectors to host cells, and that adhesins localized to the surface of these OMVs may help target effector activity to specific host tissues^37^. Similarly, vesicles, which are known to be located within mature biofilm matrices, may help to direct the bioactive OMV cargo to these conserved microcolony and EPS structures.

**Figure 6:**
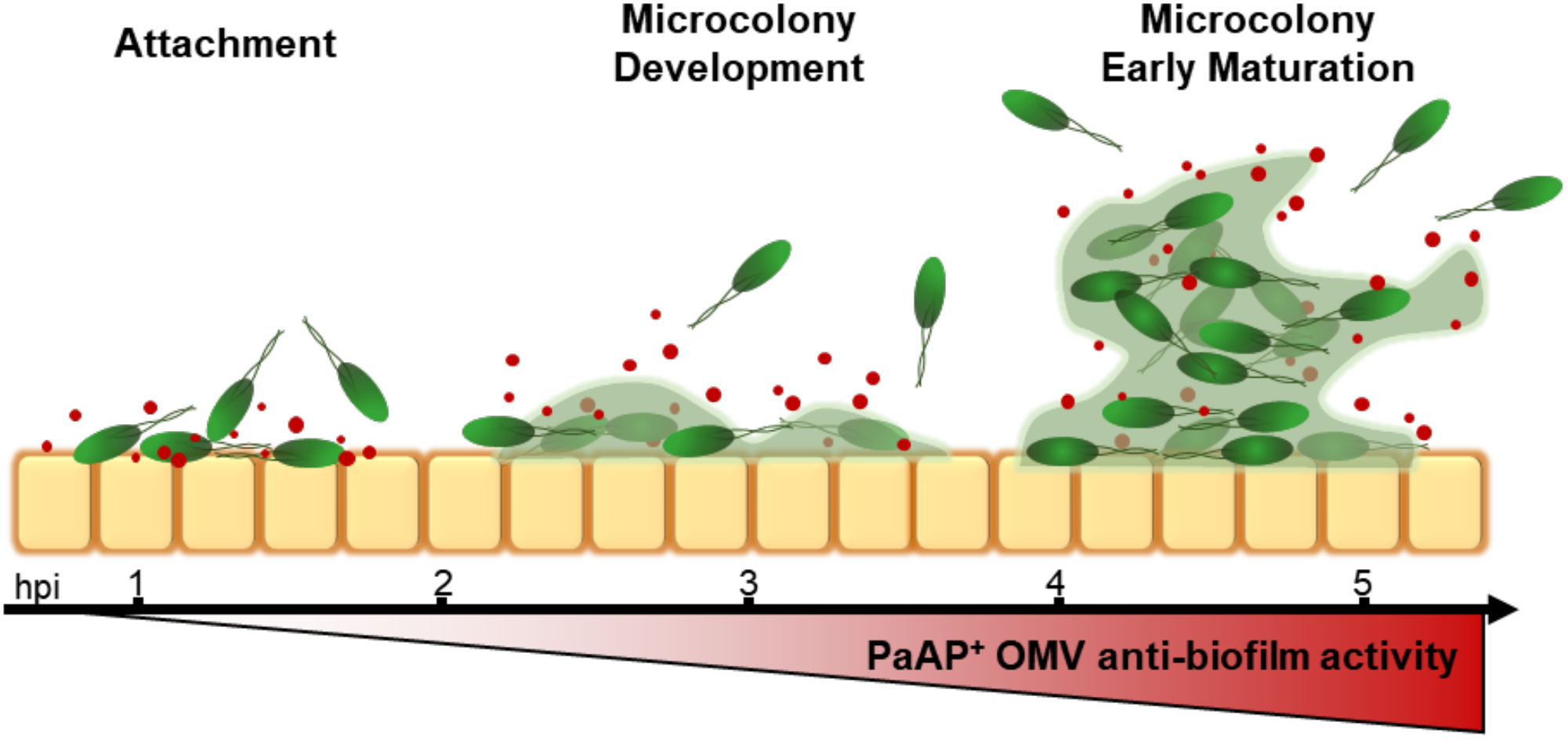
Model of PaAP^+^ OMV interaction with bacterial microcolonies and host cell substrate during biofilm coculture development stages. General stages of microcolony development are represented from left to right from time points observed in our coculture experiments (hpi: hours post infection). PaAP^+^ OMVs from the environment and those produced inside microcolonies access a conserved feature governing bacterial microcolony attachment to and development on host cells. Orange boxes: lung epithelial cells; Green ovals: *P. aeruginosa*; Red dots: OMVs; Green haze: biofilm matrix. The increasing microcolony dispersal activity of PaAP+ OMVs during the time course of microcolony development is depicted as an increase in red intensity below the figure.

Our data support the concept that increased PaAP expression observed in the clinical CF isolates is functionally similar to decreased virulence factor expression, as it limits biofilms at early microcolony formation stages, possibly to promote a less acutely toxic, long-term biofilm infection. In addition, aminopeptidase expression and secretion would allow biofilm-restricted cells to disperse and inhabit new, and perhaps more attractive, environmental niches. Typically, proteins enriched in bacterial clinical isolates contribute positively to pathogenesis mechanisms and biofilms are known to contribute to resistant infections, so PaAP’s ability to inhibit biofilm formation appears to deviate from this expectation. However, bacterial factors have been identified that can contribute to human disease in a counterintuitive fashion. Longitudinal studies that follow the progression of *P. aeruginosa* strains from early colonization through chronic infection have shown that the bacteria quickly evolve and adapt to the conditions found in the CF lung, and that many of the genetic changes seen in chronic *P. aeruginosa* strains function to decrease overall bacterial pathogenicity^32–34^. For example, *mucA* mutations, which downregulate the expression of virulence-associated type III secretion systems, are frequently observed^35,36^. Additionally, researchers have found mutations that decrease quorum sensing and, notably for this discussion, biofilm formation^32,33^. The attenuation of virulence mechanisms during infection allows pathogens such as *P. aeruginosa* the ability to thrive in the host by reducing the possibility of an overactive immune response or host cell death.

In summary, the data presented here demonstrate that deletion of the *P. aeruginosa* leucine aminopeptidase enhances early biofilm formation and microcolony development under infection-like conditions. We have also shown that this process is mediated by naturally secreted OMVs, and that exogenous addition of PaAP-containing OMVs is able to inhibit biofilm growth and disperse previously formed biofilm structures formed by both self and non-self bacterial species. Future studies will seek to determine not only the mechanism behind this activity, but also its potential clinical relevance.

## Acknowledgements

We thank David FitzGerald (National Cancer Institute, Bethesda, MD) for anti-PaAP antibody and rPaAP constructs, and acknowledge the contribution of the dTomato plasmid by Carol Kim (University of Maine, Orno ME). We acknowledge funding from NIGMS: R01GM099471-01A1, Duke University, and the NIH Cell and Molecular Biology Training Grant.

